# Genetic Structure and Diversity of Banana Bunchy Top Virus (BBTV) in the Philippines

**DOI:** 10.1101/2021.05.11.442582

**Authors:** Jay-Vee S. Mendoza, Fe M. dela Cueva, Cris Q. Cortaga, Anand Noel C. Manohar, Roanne R. Gardoce, Grace C. Lachica, Maricel C. Gonzales, John E. Thomas, Darlon V. Lantican

## Abstract

Banana bunchy top virus (BBTV) is an important disease of banana in the Philippines and in other banana-producing countries. This study was conducted to investigate the genetic structure and diversity of Philippine BBTV isolates which remain unexplored in the country. BBTV-infected plant tissues were sampled from banana-growing provinces (i.e., Cagayan, Isabela, Quirino, Batangas, Laguna, Rizal, Quezon, Palawan, Cebu, Leyte, and Davao del Sur) and the partial DNA-R gene of BBTV was sequenced. Analysis of all local BBTV isolates showed a nucleotide diversity (π) of 0.00721, average number of nucleotide differences (k) of 5.51984, and haplotype diversity (hd) of 0.971. Neutrality tests using Fu’s *Fs* and Tajima’s *D* showed significant and highly negative values which suggest an excess number of rare alleles due to recent population expansion or from genetic hitchhiking. Haplotype network and phylogenetic analyses revealed that the local BBTV isolates were closely related to Southeast Asian (SEA) group and exhibited a monophyletic clade with distinct haplotype grouping from other SEA sequences. However, some Indonesian and Indian reference sequences were also clustered within the Philippine BBTV group suggesting sequence homology. Results also showed that the local BBTV isolates may be categorized into three major haplotype groups (HA, HB, and HC) but only the HC group remained distinct upon comparison with other Philippine and SEA reference sequences. BBTV isolates from Quezon were the most diverse while isolates from Palawan displayed low genetic diversity indices and belonged only in the HC group. The assessment of the degree of variability among Philippine BBTV isolates will provide a reference towards the development of high-throughput BBTV detection systems as well as enable to devise plant breeding strategies to manage the current BBTV spread and variations.

## Introduction

Banana bunchy top disease (BBTD) is one of the major threats in banana producing countries such as India, China, Taiwan, Indonesia and the Philippines (Debbarma et al., 2019; Qazi, 2016). In the Philippines, BBTD is present in almost all banana-growing areas. The disease affects both smallholders and large banana plantation growers. Spread of the disease can be minimized through elimination of infected materials and use of virus-free tissue culture-derived planting materials. These management strategies are usually employed by big commercial growers instead of the smallholders. In the 1990s, smallholder production of the popular dessert cv. Lakatan in the northern Philippines was virtually eliminated by severe BBTD infection (Molina et al., 2009). Initial symptoms of the disease involve the appearance of dark green streaks in the veins which becomes apparent with a combination of marginal chlorosis or yellowing of the leaves. Dashes and dots creating a “Morse code” pattern in the leaves and petiole may also be observed. This will lead into small emerging bunchy leaves forming a rosette pattern. In serious cases, disease plants were observed to be severely stunted and unable to bear fruits (Dale 1987; Hooks et al., 2008).

The disease is known to be caused by Banana bunchy top virus (BBTV), a single-stranded DNA virus belonging to the Babuvirus genus, which can bring catastrophic loss to a banana plantation. The systemic virus can easily be transmitted by an aphid (*Pentalonia nigronervosa*; Magee, 1927), a vector with a wide host range including *Musa textilis* and other members of the family Musaceae. The disease transmission is of the persistent, circulative, non-propagative type (Anhalt and Almeida, 2008), with efficiency ranging from 46-67% (Magee, 1927; Wu and Su., 1990; Hu et al., 1996). The virus is made up of 6 genetic components, namely, DNA-C coding for the cell cycle link protein, DNA-S coding for the capsid protein, DNA-M coding for the movement protein, DNA-N coding for the nuclear shuttle protein, DNA-U3 coding for potential protein with unknown function, and DNA-R coding for the replication initiation protein (Amin et al., 2008, Kumar et al., 2017; Wickramaarachchi et al., 2016).

The phylogenetic relationship among BBTV DNA-R sequences revealed that the virus can be categorized into two different lineages based on geographical distribution: the South Pacific/ Pacific-Indian Oceans (PIO) group and the Asian/ Southeast Asian (SEA) group (Yu et al., 2012; Karan et al., 1994). BBTV isolates obtained from Australia, Egypt, Hawaii, India, Myanmar, Pakistan, Sri Lanka and Tonga belong under the PIO group, while the isolates collected from China, Indonesia, Japan, Philippines, Taiwan and Vietnam are considered members of the SEA group (Yu et al., 2012).

In-depth studies on the diversity of BBTV from various countries were already reported, such as in Democratic Republic of Congo (Mukwa et al., 2016), Pakistan (Amin et al., 2007), sub-Saharan Africa (Kumar et al., 2011), Indonesia (Chiaki et al., 2015), Japan (Furuya et al., 2005) and India (Banerjee et al., 2014). The information on the degree of genetic diversity and distribution of BBTV in these countries provided useful and fundamental information to control BBTD through various pest management approaches. Unfortunately, the reported BBTV sequences from the Philippines have only been made available through foreign efforts, which generally aims to provide insights into BBTV diversity and population structure at the global level. Local and intensive reports regarding the diversity and spread of this important banana disease will be vital towards devising a specific management system in the country. Thus, a detailed assessment of the current Philippine BBTV diversity and genetic structure using the DNA-R region will be discussed in this paper.

## Materials and Methods

### Survey and collection

A survey was done in banana growing areas in the Philippines (Figure 1). Symptoms depicting classic BBTV infection such as leaf chlorosis, dash-dot pattern, rosetting, and stunting were observed and recorded. Leaf samples from representative symptomatic and asymptomatic samples were collected and processed for DNA extraction.

**Figure 1.**
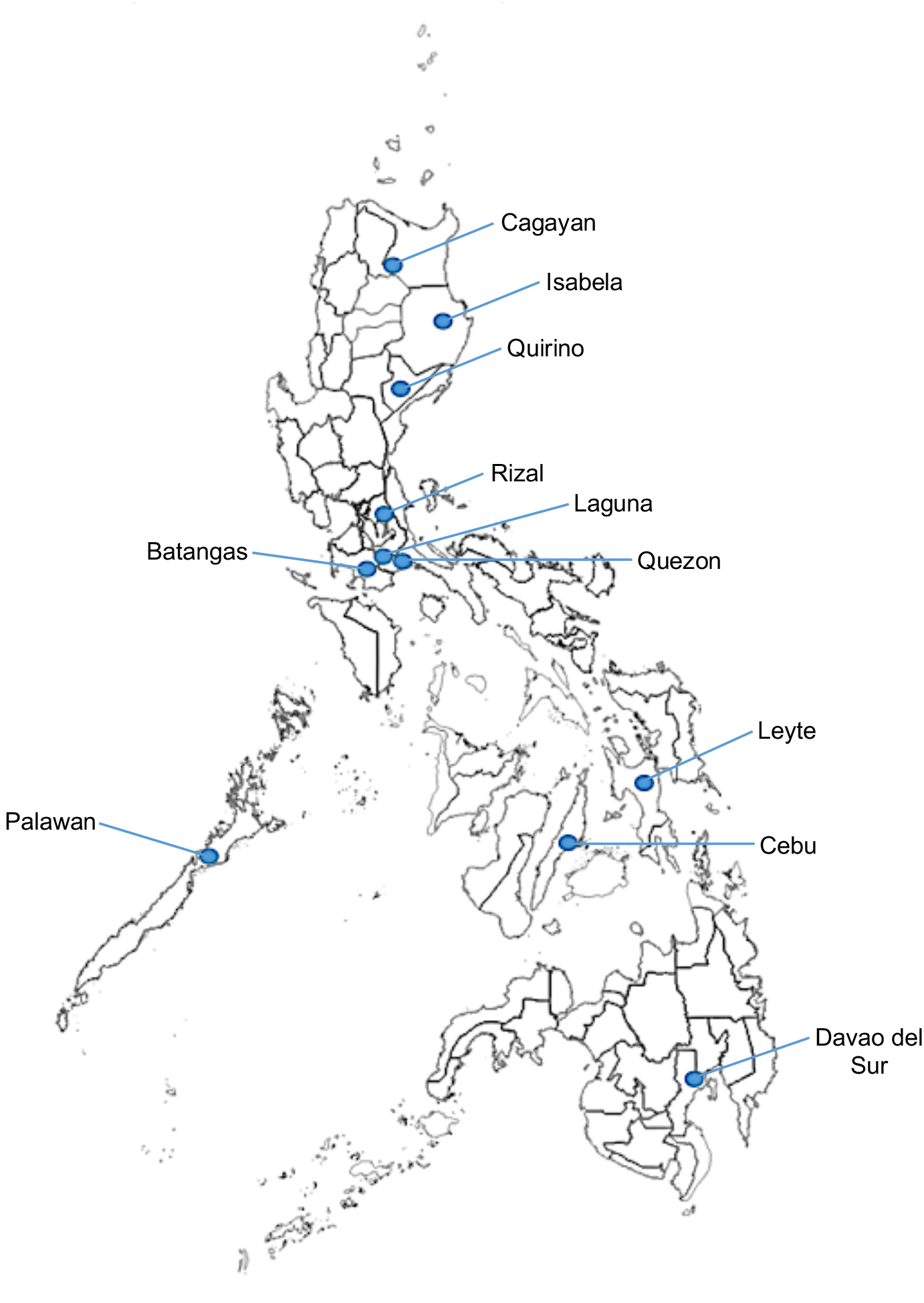
Geographical distribution of BBTD collected from banana-growing areas in the Philippines.

### DNA extraction

Samples were processed using a CTAB DNA extraction protocol adapted from Doyle and Doyle (1990) with a few modifications. Approximately, 300 mg of fresh leaf tissue was ground into fine powder with liquid nitrogen using a sterilized mortar and pestle. Ground tissue was then transferred into a sterilized 1.5 ml microcentrifuge tube. Exactly 700 µl of extraction buffer with 2% PVP was added to each sample and then incubated at 65 °C for an hour. One volume CH_3_Cl-isoamyl alcohol (24:1) was added followed by centrifugation at 10,000 rpm for 10 minutes at 23°C. Aqueous phase was transferred into a new sterile 1.5 ml microcentrifuge tube. DNA was precipitated by adding 0.8 volume of cold isopropanol and incubated at -20 °C for 30 minutes, followed by centrifugation at 10,000 rpm for 15 minutes. The DNA pellet was washed using 1 mL of Wash 1 (0.2 M sodium acetate, 76% ethanol; filter sterilized) for 10 minutes followed by 1 ml of Wash 2 (10 mM ammonium acetate, 76% ethanol; filter sterilized) for 5 minutes, and the pellet was air-dried for 30 minutes. The DNA pellet was resuspended in Tris-EDTA buffer (10 mM Tris-HCl, 1 mM disodium EDTA, pH 8.0) and purified by incubation with 0.1 mg/ml RNase at 37 °C for 1 hour, and centrifugation at 10,000 rpm for 5 minutes. DNA was collected into individual sterile 1.5 ml microcentrifuge tubes and stored at -20 °C.

### PCR detection

The presence of BBTV was confirmed by performing PCR detection. Each 15 µL reaction mixture is consisting of 1X PCR buffer (10 mM Tris pH 9.1 at 20 °C, 50 mM KCl, 0.01% Triton™ X-100; Vivantis Technologies, Malaysia), 1.76 mM MgCl_2_, 0.2mM dNTPs, 2 µM of BBT1 (5’-CTC GTC ATG TGC AAG GTT ATG TCG-3’) and BBT2 (5’-GAA GTT CTC CAG CTA TTC ATC GCC-3’) primers (Thompson and Dietzgen, 1995; Harding et al., 1993; Integrated DNA Technologies Pte. Ltd., Singapore), 1U of Taq Polymerase (Vivantis Technologies, Malaysia), and 20 ng DNA. The PCR mixture was then run in a T100 thermal cycler (BioRad, USA) with initial denaturation at 94 °C for 10 minutes, followed by 30 cycles of 94 °C for 1 minute, 53 C for 1 minute, 72 °C for 2 minutes and a final extension of 72 °C for 10 minutes. PCR products were viewed with electrophoresis using 1% agarose gels in 1X TBE buffer at 100 V for 40 min and visualized using 0.5 ug/ml ethidium bromide staining and UV illumination using the Enduro GDS Touch Imaging System (Labnet International, Inc, Edison, New Jersey, USA).

### Outsourced sequencing

Confirmed BBTV isolates were further processed for the partial sequencing of the DNA-R region of the virus (Table 1). PCR amplification was done using previously established primers for assessing genetic similarity (Islam et al., 2010): BBTVREP-F (5’-ATG GCG CGA TAT GTG GTA TGC -3’) and BBTVREP-R (5’-TCA GCA AGA AAC CAA CTT TAT TCG - 3’). The DNA-R primer was optimized using the same PCR conditions as BBT1 and BBT2 primers. PCR products were then sent for outsourced capillary sequencing (Apical Scientific, Malaysia).

**Table 1.** Collected BBTV Philippine isolates sequenced in this study.

| Isolate | Location | Collection Year | Host variety |
| --- | --- | --- | --- |
| Balam_Cebu_B22 | Balamban, Cebu | 2015 | Unknown |
| Baybay_Ley_B34 | Baybay, Leyte | 2015 | Unknown |
| Baybay_Ley_B35 | Baybay, Leyte | 2015 | Unknown |
| BenSol_Isab_B18 | Benin Soledad, Isabela | 2015 | Unknown |
| Calin_Dav_B12 | Calinan, Davao City | 2015 | Unknown |
| Calin_Dav_B14 | Calinan, Davao City | 2015 | Unknown |
| Can_Que3 | Candelaria, Quezon | 2019 | Unknown |
| Can_Que_07 | Candelaria, Quezon | 2019 | Tordan and Saba |
| Can_Que_08 | Candelaria, Quezon | 2019 | Tordan and Saba |
| Can_Que_11 | Candelaria, Quezon | 2019 | Tordan and Saba |
| Can_Que_12 | Candelaria, Quezon | 2019 | Tordan and Saba |
| Can_Que13 | Candelaria, Quezon | 2019 | Tordan and Saba |
| Can_Que15 | Candelaria, Quezon | 2019 | Tordan and Saba |
| Diffun_Qui_B19 | Diffun, Quirino | 2015 | Unknown |
| Luc_Que_01 | Lucban, Quezon | 2019 | Unknown |
| Luc_Que_02 | Lucban, Quezon | 2019 | Unknown |
| Luc_Que_05 | Lucban, Quezon | 2019 | Lakatan Tagalog |
| Luc_Que4 | Lucban, Quezon | 2019 | Lakatan Tagalog |
| Luc_Que10 | Lucban, Quezon | 2019 | Lakatan Tagalog |
| Luc_Que6 | Lucban, Quezon | 2019 | Lakatan Tagalog |
| Luis_Lag_04 | Luisiana, Laguna | 2019 | Lakatan |
| Luis_Lag_07 | Luisiana, Laguna | 2019 | Lakatan |
| Luis_Lag_08 | Luisiana, Laguna | 2019 | Lakatan |
| Luis_Lag6 | Luisiana, Laguna | 2019 | Lakatan |
| Luis_Lag7 | Luisiana, Laguna | 2019 | Lakatan |
| Magda_Lag_B10 | Magdalena, Laguna | 2015 | Unknown |
| Naga_Cebu_B24 | Naga, Cebu | 2015 | Unknown |
| Orm_Ley_B31 | Ormoc, Leyte | 2015 | Unknown |
| Orm_Ley_B32 | Ormoc, Leyte | 2015 | Unknown |
| Orm_Ley_B32b | Ormoc, Leyte | 2015 | Unknown |
| Pagsan_Lag_01 | Pagsanjan, Laguna | 2019 | Saba |
| Pagsan_Laguna2 | Pagsanjan, Laguna | 2019 | Saba |
| Rizal_Cag_B16 | Rizal, Cagayan | 2015 | Unknown |
| Roxas_Pal_01 | Roxas, Palawan | 2019 | Lakatan and Saba |
| Roxas_Pal_03 | Roxas, Palawan | 2019 | Lakatan and Saba |
| Roxas_Pal_04 | Roxas, Palawan | 2019 | Lakatan and Saba |
| Roxas_Pal_05 | Roxas, Palawan | 2019 | Lakatan and Saba |
| Roxas_Pal_07 | Roxas, Palawan | 2019 | Lakatan |
| Roxas_Pal12 | Roxas, Palawan | 2019 | Lakatan |
| Roxas_Pal15 | Roxas, Palawan | 2019 | Lakatan |
| Roxas_16 | Roxas, Palawan | 2019 | Lakatan |
| Roxas_Pal2 | Roxas, Palawan | 2019 | Lakatan |
| Roxas_Pal5 | Roxas, Palawan | 2019 | Lakatan |
| Talisay_Bat_08 | Talisay, Batangas | 2019 | Saba |
| Talisay_Bat_09 | Talisay, Batangas | 2019 | Saba |
| Talisay_Bat_10 | Talisay, Batangas | 2019 | Saba |
| Talisay_Bat_10b | Talisay, Batangas | 2019 | Saba |
| Talisay_Bat3 | Talisay, Batangas | 2019 | Saba |
| Talisay_Bat4 | Talisay, Batangas | 2019 | Saba |
| Talisay_Bat6 | Talisay, Batangas | 2019 | Saba |
| Tanauan_Bat_11 | Tanauan, Batangas | 2019 | Latundan |
| Tanauan_Bat_16 | Tanauan, Batangas | 2019 | Latundan |
| Tanauan_Bat_17 | Tanauan, Batangas | 2019 | Latundan |
| Tanauan_Bat_18 | Tanauan, Batangas | 2019 | Latundan |
| Tanauan_Bat_20 | Tanauan, Batangas | 2019 | Latundan |
| Tanauan_Bat19 | Tanauan, Batangas | 2019 | Latundan |
| Tanauan_Bat11 | Tanauan, Batangas | 2019 | Latundan |
| Tanay_Riz_B01 | Tanay, Rizal | 2015 | Unknown |
| Tanay_Riz_B02 | Tanay, Rizal | 2015 | Unknown |
| Tanay_Riz_B03 | Tanay, Rizal | 2015 | Unknown |
| Tanay_Riz_B04 | Tanay, Rizal | 2015 | Unknown |
| Tugbok_Dav_B11 | Tugbok, Davao City | 2015 | Unknown |
| Tugbok_Dav_B13 | Tugbok, Davao City | 2015 | Unknown |
| Tugbok_Dav_B15 | Tugbok, Davao City | 2015 | Unknown |

### Sequence analysis

Raw paired sequences (forward and reverse) were quality trimmed and analyzed using Geneious Prime® (version 2019.0.4). Trimmed and assembled sequences were then aligned using ClustalW (Thompson et al., 2003) at default settings. Resulting alignment was then used for phylogenetic analysis. The partial DNA-R sequences from collected BBTV samples were compared with published reference sequences in NCBI (Appendix Table 1) to determine its relation to the South Pacific group and Asian group and to confirm the identity of the virus. Two (2) Abaca bunchy top virus (ABTV) reference sequences (accession numbers: EF546813.1, EF546807.1) served as outgroups in the analysis.

### Genetic diversity and demographic analyses

Parameters of genetic diversity and demographic analysis using the partial DNA-R gene of BBTV populations isolated in the Philippines were computed using DNA Sequence Polymorphism (DnaSP) (Rozas et al., 2017). The estimates of evolutionary divergence (genetic distance) over sequence pairs between and within population (inter- and intra-population, respectively) of BBTV isolates were computed using Molecular Evolutionary Genetics Analysis (MEGA X) (Kumar et al., 2018) based on T93 nucleotide substitution model (Tamura and Nei 1993). The rate variation among sites was modeled with a gamma distribution (+G) and discreet evolutionary invariable sites (+I).

### Haplotype network and phylogenetic analyses

Based on DNA-R sequence of BBTV, two haplotype networks were constructed using Population Analysis with Reticulate Trees (PopART) (Leigh and Bryant, 2015): (1) median joining network of collected Philippine isolates; and (2) minimum spanning network of collected Philippine isolates with other published reference sequences from the Philippines and SEA. The BBTV phylogenetic tree was reconstructed using the maximum likelihood statistical method implemented in IQ-TREE (Nguyen et al., 2015) with best-fit substitution model selected based on Bayesian information criterion (BIC) through ModelFinder (Kalyaanamoorthy et al., 2017). The tree was generated using TIM2 model (AC=AT, CG=GT and unequal base frequency; Posada 2008) with empirical base frequencies (+F) and FreeRate heterogeneity across sites model (+R3) (Yang, 1995; Soubrier et al., 2012). The resulting phylogenetic tree was validated with 1,000 replicates of ultrafast bootstrapping (Hoang et al., 2018) and visualized using FigTree (Rambaut, 2018).

## Results

### Genetic diversity and demographic analysis

Banana leaves showing characteristic symptoms of banana bunchy top disease (BBTD) were collected from 11 banana growing areas in the country, namely, Cagayan, Isabela, Quirino, Batangas, Laguna, Rizal, Quezon, Palawan, Cebu, Leyte, and Davao del Sur (Figure 1). Among the BBTV populations with partial DNA-R sequences (Table 1), the highest number of segregating sites (S) was observed in Quezon and Batangas (S=25 and 28, respectively) while the lowest was observed in Rizal and Cebu (S=6) (Table 2). The nucleotide diversity (π) was highest in Quezon (π=0.00893) and lowest in Rizal (π=0.00385). The average number of nucleotide differences (k) was highest in Quezon, Cebu, and Leyte (ranging from 6 to 6.897) and lowest in Rizal, Palawan, and Leyte (ranging from 3 to 3.2). The number of haplotype (h) was highest in Batangas and Quezon (h=12) with relatively high haplotype diversity (hd) of 0.967 and 0.987, respectively. A high haplotype diversity (hd) of 1 was observed in Cebu (h=2), Laguna (h=8), and Davao del Sur (h=5) as the haplotype number in these locations corresponded to the number of samples obtained. Lowest haplotype diversity (hd) was observed in Palawan (0.644). Analysis of all Philippine BBTV populations showed 59 total segregating sites (S) with nucleotide diversity (π) of 0.0021 and average number of nucleotide differences (k) of 5.51984, while the haplotype number (h) was 41 with haplotype diversity (hd) of 0.971.

**Table 2.** Parameters of genetic diversity and demographic analysis using DNA-R gene of BBTV populations isolated in the Philippines.

| Location | n | S | | | $\pi$ (SD) | k | h | hd (SD) | Fu's *F<sub>s</sub>* | | Tajima's *D* | |
| --- | --- | --- | --- | --- | --- | --- | --- | --- | --- | --- | --- | --- |
|  |  | SS | PIS | Total S |  |  |  |  | *F<sub>s</sub>* | *P* value<sup>a</sup> | *D* | Significance |
| **Cagayan** | 1 | - | - | - | - | - | - | - | - | - | - | - |
| **Isabela** | 1 | - | - | - | - | - | - | - | - | - | - | - |
| **Quirino** | 1 | - | - | - | - | - | - | - | - | - | - | - |
| **Batangas** | 14 | 23 | 5 | 28 | 0.00653 (0.00133) | 5.07692 | 12 | 0.967 (0.044) | -5.478 | 0.00400 | -1.80970 | *P* < 0.05* |
| **Laguna** | 8 | 7 | 5 | 12 | 0.00567 (0.00057) | 4.39286 | 8 | 1.000 (0.063) | -4.309 | 0.00602 | -0.25574 | *P* > 0.10<sup>ns</sup> |
| **Rizal** | 4 | 6 | 0 | 6 | 0.00385 (0.00167) | 3.00000 | 3 | 0.833 (0.222) | 0.731 | 0.62381 | -0.80861 | *P* > 0.10<sup>ns</sup> |
| **Quezon** | 13 | 15 | 10 | 25 | 0.00893 (0.00162) | 6.89744 | 12 | 0.987 (0.035) | -5.026 | 0.01200 | -0.90389 | *P* > 0.10<sup>ns</sup> |
| **Palawan** | 10 | 8 | 3 | 11 | 0.00416 (0.00158) | 3.22222 | 3 | 0.644 (0.101) | 3.321 | 0.94790 | -0.76710 | *P* > 0.10<sup>ns</sup> |
| **Cebu** | 2 | 6 | 0 | 6 | 0.00772 (0.00386) | 6.00000 | 2 | 1.000 (0.500) | 1.792 | 0.53944 | - | - |
| **Leyte** | 5 | 8 | 0 | 8 | 0.00411 (0.00162) | 3.20000 | 3 | 0.700 (0.218) | 1.458 | 0.75358 | -1.17432 | *P* > 0.10<sup>ns</sup> |
| **Davao del Sur** | 5 | 5 | 7 | 12 | 0.00798 (0.00164) | 6.20000 | 5 | 1.000 (0.126) | -1.011 | 0.15015 | 0.55247 | *P* > 0.10<sup>ns</sup> |
| **ALL** | 64 | 37 | 22 | 59 | 0.00721 (0.00053) | 5.51984 | 41 | 0.971 (0.011) | -33.210 | 0.00000 | -1.98369 | *P* < 0.05* |

n=no. of BBTV sequences; S=segregating sites; SS=singleton sites; PIS=parsimony informative sites; $\pi$ =nucleotide diversity; k=average no. of nucleotide differences, h=no. of haplotypes; hd=haplotype diversity; SD=standard deviation; \*=significant; <sup>a</sup> should be regarded as significant (5% level) if *P* < 0.02.

For the test of neutrality (Table 2), significantly (*P* < 0.02) negative Fu’s *Fs* was observed in BBTV population from Batangas (−5.478), Laguna (−4.309), and Quezon (−5.026) (Table 2), while only Batangas has significantly (*P* < 0.05) negative Tajima’s *D* (−1.80970). Meanwhile, Tajima’s *D* was not computed for Cebu samples due to small sample size. Analysis of all Philippine BBTV isolates showed a significant and highly negative Fu’s *Fs* (−33.210) and Tajima’s D (−1.98369).

BBTV isolates from Cebu, Leyte, Davao del Sur, Quezon, Laguna, and Batangas showed the highest inter-population genetic distance with Isabela isolates (ranging from 0.009 to 0.011), while BBTV isolates from Quirino, Cagayan, Palawan, and Rizal showed the highest inter-population genetic distance with Batangas isolates (ranging from 0.007 to 0.010) (Table 3). Intra-population genetic distance was highest in Quezon isolates, while Rizal isolates showed the lowest.

**Table 3.** Estimates of evolutionary divergence (inter- and intra-population genetic distance) of BBTV populations collected in the Philippines.

| Location | Inter-population genetic distance |  |  |  |  |  |  |  |  |  | Intra-population genetic distance |  |
| --- | --- | --- | --- | --- | --- | --- | --- | --- | --- | --- | --- | --- |
|  | Cebu | Leyte | Isabela | Davao del Sur | Quezon | Quirino | Laguna | Cagayan | Palawan | Batangas | Distance | SE |
| **Cebu** |  |  |  |  |  |  |  |  |  |  | 0.00513 | 0.00259 |
| **Leyte** | 0.00644 |  |  |  |  |  |  |  |  |  | 0.00360 | 0.00142 |
| **Isabela** | 0.01037 | 0.01011 |  |  |  |  |  |  |  |  | - | - |
| **Davao del Sur** | 0.00592 | 0.00618 | 0.00959 |  |  |  |  |  |  |  | 0.00566 | 0.00196 |
| **Quezon** | 0.00700 | 0.00769 | 0.01017 | 0.00705 |  |  |  |  |  |  | 0.00809 | 0.00201 |
| **Quirino** | 0.00644 | 0.00618 | 0.00385 | 0.00566 | 0.00645 |  |  |  |  |  | - | - |
| **Laguna** | 0.00499 | 0.00628 | 0.01020 | 0.00596 | 0.00711 | 0.00628 |  |  |  |  | 0.00392 | 0.00140 |
| **Cagayan** | 0.00514 | 0.00489 | 0.00516 | 0.00437 | 0.00555 | 0.00128 | 0.00627 |  |  |  | - | - |
| **Palawan** | 0.00789 | 0.00763 | 0.00790 | 0.00680 | 0.00769 | 0.00400 | 0.00885 | 0.00271 |  |  | 0.00370 | 0.00126 |
| **Batangas** | 0.00707 | 0.00865 | 0.01112 | 0.00781 | 0.00838 | 0.00737 | 0.00570 | 0.00831 | 0.01022 |  | 0.00619 | 0.00143 |
| **Rizal** | 0.00595 | 0.00534 | 0.00614 | 0.00489 | 0.00613 | 0.00225 | 0.00676 | 0.00096 | 0.00368 | 0.00894 | 0.00192 | 0.00107 |
| **Mean distance (SE): 0.00722 (0.00162)** |  |  |  |  |  |  |  |  |  |  |  |  |

SE=standard error

The data on genetic diversity, demographic analysis, and intra-population genetic distance were not computed for BBTV isolates from Cagayan, Isabela, and Quirino as only one representative isolate was obtained from each location.

### Haplotype analysis

Haplotype network of BBTV samples isolated from the Philippines revealed three major haplotype groups (HA, HB, and HC) using the partial DNA-R gene (Figure 2a). The first haplotype group (HA) includes isolates from Laguna, Batangas, and Quezon which are provinces from the Luzon region. The second haplotype group (HB) includes isolates from Laguna and Quezon (Luzon region), Cebu and Leyte (Visayas region), and Davao del Sur (Mindanao region). The third haplotype group (HC) includes isolates from all sampled provinces (except Laguna and Batangas). Interestingly, all Palawan isolates were included in this haplotype group only. Additional BBTV samples and genes will be analyzed to confirm the haplotype groupings observed.

**Figure 2.**
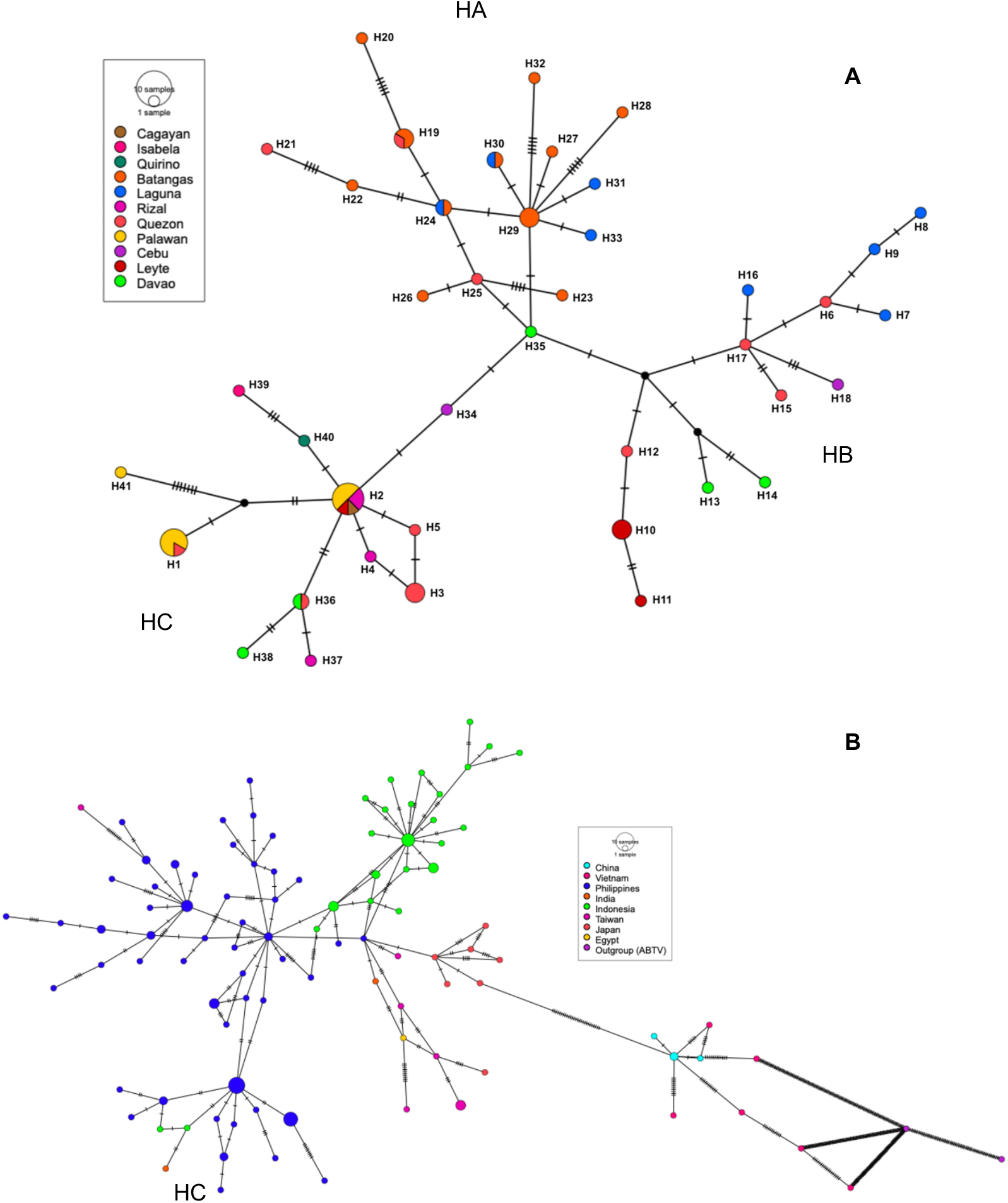
Haplotype network based on partial DNA-R sequence of BBTV constructed using PopART (Leigh and Bryant, 2015): median joining network of the collected Philippine isolates (A) and minimum spanning network of collected isolates with Philippine and other SEA reference sequences (B). Haplotype Groups: HA, HB, HC.

Using the DNA-R gene, a SEA haplotype network consisting of collected Philippine isolates and reference sequences from the Philippines and SEA was constructed (Figure 2b). It showed distinct grouping of Philippine isolates from its neighboring Asian countries (such as Indonesia, Taiwan, Japan), while China and Vietnam showed the most distant haplotype grouping. Here, the HC group remains distinct wherein BBTV sequences from Indonesia (n=2) and India (n=1) were also included.

### Phylogenetic analysis

Phylogenetic analysis showed two broad clades/groups of BBTV, namely, the SEA and PIO groups, with high bootstrap support values of 88% and 90%, respectively (Figure 3). All Philippine BBTV sequences were found in the SEA clade. The Philippine reference sequences and Philippine BBTV isolates in this study clustered together wherein sequences from Indonesia (n=8) and India (n=1) were also included. BBTV sequences from India and Egypt were both found in SEA and PIO clades. Notably, the clustering of Philippine BBTV isolates appeared to follow the three haplotype groupings observed in this study (HA, HB, and HC). Philippine reference sequences were found among the HA and HB isolates but not on HC isolates. On the other hand, Indonesian sequences were found among HB and HC isolates, while an Indian BBTV sequence was found among HC isolates. In the SEA clade, China and Vietnam sequences formed a separate cluster from the rest of SEA sequences with a well-supported bootstrap value of 82%.

**Figure 3.**
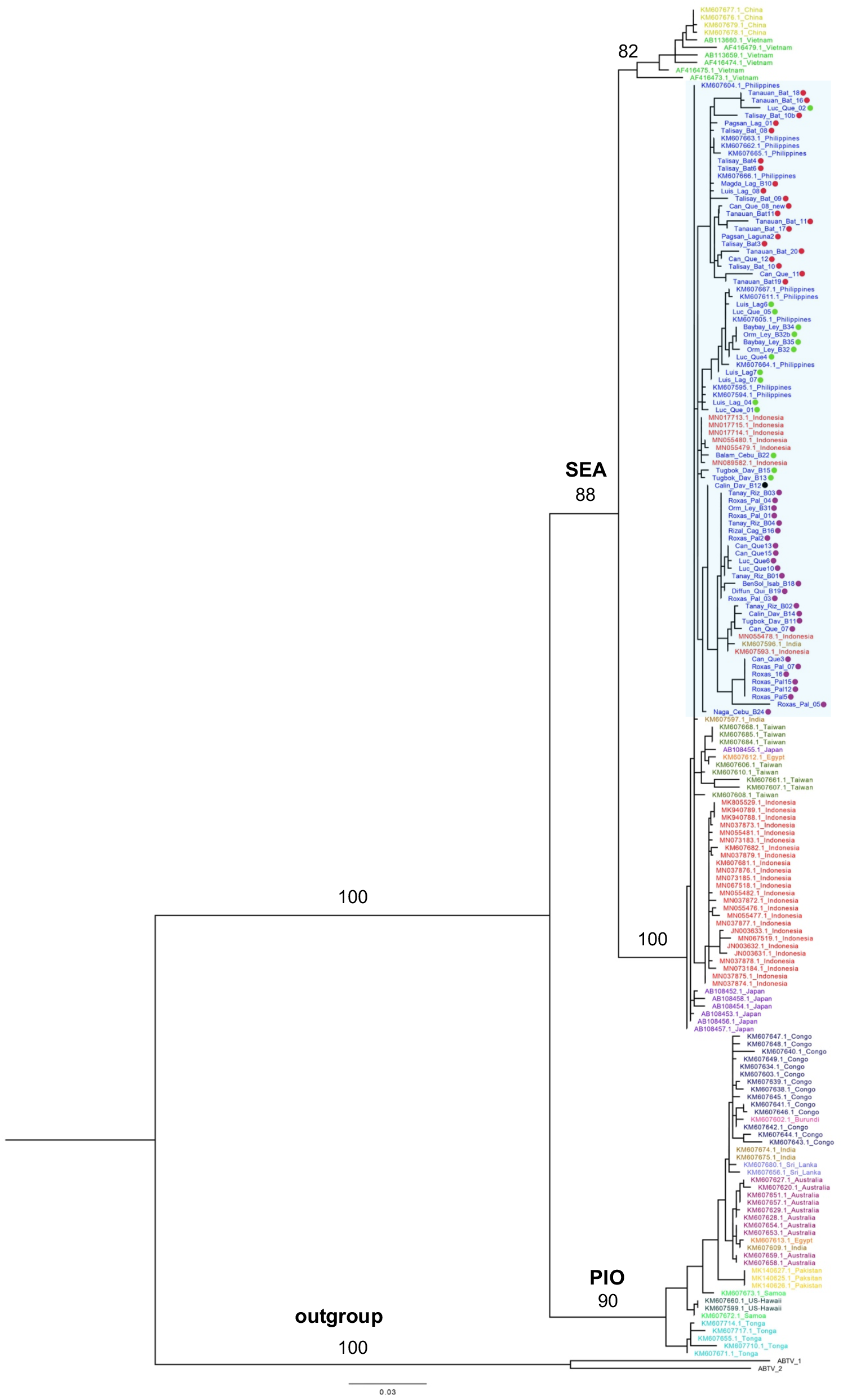
Maximum likelihood phylogenetic tree constructed using IQ-TREE from the partial DNA-R sequence alignment of collected Philippine BBTV isolates and published Philippine, SEA, and PIO reference sequences. Best-fit model was selected according to BIC using ModelFinder (Kalyaanamoorthy et al., 2017). The tree was generated using TIM 2 model (AC=AT, CG=GT and unequal base frequency; Posada 2008) with empirical base frequencies (+F) and FreeRate heterogeneity across sites model (+R3) (Yang, 1995; Soubrier et al., 2012). The tree was tested with 1,000 replicates of ultrafast bootstrapping (Hoang et al., 2018) and visualized using FigTree (v1.4.4) (Rambaut 2018). The numbers in the branches are bootstrap support values. Clade highlighted in blue contains the collected Philippine BBTV isolates with colored dots corresponding to haplotype grouping: red dot = HA, green dot = HB, and purple dot = HC. SEA = Southeast Asian group, PIO = Pacific-Indian Oceans group. ABTV sequences served as outgroups.

## Discussion

Banana bunchy top disease has been an important disease of the banana crop in the Philippines (Molina et al., 2009). Molecular information regarding the Philippine BBTV isolates has been lacking and remains unexplored. Thus, this study was performed to investigate the genetic structure and diversity of the BBTV isolates in the Philippines. Here, the partial DNA-R region of BBTV was sequenced due to its wide application in assessing genetic diversity and other molecular analyses (Bell et al., 2002; Furuya et al., 2005; Amin et al., 2007; Kumar et al., 2011; Shekhawat et al., 2012; Banerjee et al., 2014; Chiaki et al., 2015; Mukwa et al., 2016). Survey and sample collection were conducted in banana growing areas in various regions in the Philippines for BBTV detection and diversity analysis. The banana cultivars wherein BBTV was isolated include Saba, Lakatan, Latundan, and other unknown varieties.

Among the sampling sites, Quezon appear to have the most diverse BBTV population as shown by high nucleotide segregating sites (S), nucleotide diversity (π), average number of nucleotide differences (k), haplotype number (h), haplotype diversity (hd), intra-population genetic distance, and a significant negative Fu’s *Fs* value. Significant and negative Fu’s *Fs* was observed also in Laguna BBTV population, while significant and negative Fu’s *Fs* and Tajima’s *D* values were observed in BBTV population from Batangas. Fu’s *Fs* is regarded as a more sensitive indicator of population expansion and a more powerful test of neutrality than the Tajima’s *D*, which probably contributed to the inconsistent results (Zeng et al., 2006). More BBTV samples should be collected from different provinces for genetic diversity and demographic analyses to confirm the results obtained. Overall, significant and highly negative Fu’s *Fs* and Tajima’s *D* were observed using all Philippine BBTV isolates. These results suggest that there is an excess number of rare alleles in Philippine BBTV isolates, probably due to its recent population expansion (or from genetic hitchhiking) as evidenced also by overall high haplotype diversity with relatively low overall nucleotide diversity.

Haplotype network and phylogenetic analyses of partial DNA-R of combined SEA and Philippine sequences suggest that geographic location heavily affects the distribution of BBTV as indicated by geographically proximate haplotypes in each group. Viruses may have evolved independently mainly because countries are separated by sea, and host movement could have been limited within the haplotype groupings. The complex haplotype network of BBTV isolates suggests that the Philippines, as part of SEA, is a hotspot of an on-going BBTV diversification (Stainton et al., 2015). Furthermore, the Philippines, along with other neighboring countries such as New Guinea and Indonesia, are believed to be the center of origin of domesticated bananas (Perrier et al., 2011). This may indicate that the intensive domestication of bananas within the region might have been a possible driver for the diversification of BBTV in the country. Meanwhile, the haplotype network analysis revealed three major haplotype groups (HA, HB, and HC) of BBTV isolates collected in the Philippines. Interestingly, BBTV isolates from Palawan were only found in the HC group. This province also has very low haplotype number (h) and haplotype diversity (hd), and a relatively low nucleotide diversity (π). These results could be probably caused by recent population bottleneck and recent introduction of BBTV in the area due to the movement of planting materials (e.g., Lakatan variety) from other provinces in the Philippines. Due to the strict quarantine implementation in the province, BBTV has been reported only recently in Palawan. It appears that the HC group is widespread in the country and was also introduced in Palawan. Upon inclusion of Philippine and SEA reference sequences in the haplotype network, however, only the HC group remained distinct wherein few BBTV reference sequences from India and Indonesia were also included.

As expected, phylogenetic analysis based on partial DNA-R showed that the collected local isolates were more closely related with the SEA group (where the Philippines is geographically classified) than the PIO group (Karan et al., 1994). This could also mean that plant and virus movement is limited within the SEA region (Karan et al., 1994; Wickramaarachchi 2016). Philippine BBTV isolates formed a monophyletic clade which suggests a monophyletic origin of the majority of local isolates from a common SEA ancestor. The collected local isolates also clustered with Philippine reference sequences which confirms their identity as BBTV and may indicate that virus movement could be limited in the country (Stainton et al., 2015). In the phylogenetic tree, the clustering of Philippine BBTV isolates seemed to follow the observed three haplotype groupings (HA, HB, and HC). However, as shown in the tree, no Philippine reference sequences appeared to cluster with HC isolates. A more exhaustive survey of reference sequences in the Philippines will be performed and more BBTV samples in the Philippines will be sequenced to verify the findings on haplotype groupings and phylogenetic analysis.

Some BBTV sequences from Indonesia and India were also clustered within the Philippine clade (the former clustered with HB and HC isolates, while the latter clustered with HC isolates), suggestive of sequence homology and possible BBTV movement in these countries. It was previously inferred that the Indian subcontinent is a major contributor to the long-distance dispersal of BBTV, both as donor and recipient. For instance, the introduction events of SEA isolates were recently detected between 1976 and 1991 in India (Stainton et al., 2015). Thus, it may be deduced that there is a probable dispersal event of Philippine BBTV isolates to India. Outside the Philippine clade, BBTV sequences from Egypt and India (which are known to be closely related with PIO group) were also clustered in the larger SEA clade, indicating the presence of isolates that are related with SEA group. On the other hand, reference sequences from Vietnam and China formed a monophyletic clade and appeared to be separated from the larger SEA group. Similar observation was reported by Rao (2017) wherein the constructed DNA-R phylogenetic tree depicts a further separation of BBTV isolates from China and Vietnam into sub-groups 2, 3, and 4; while the rest of the members of the Asian group exclusively formed the sub-group 1.

In summary, the results of this study showed that BBTV is widespread and diverse in the Philippines and undergoing population expansion. However, more samples and genes should be analyzed to confirm the results obtained especially at the province level. Additional reference sequences from the Philippines and other countries with reported BBTV occurrence will be included in the analysis. Recombination analysis shall be also performed to provide further understanding regarding the evolutionary history of Philippine BBTV isolates. Nevertheless, the insights drawn from this research endeavor will provide a framework in the development of improved BBTV-specific detection marker systems in the country as well enable the strategic BBTV-resistant variety deployment across various regions in the Philippines.

## Acknowledgment

This research was conducted through funds provided by Philippine Department of Agriculture - Biotechnology Program Office (DA-BIOTECH) to the project entitled “DA-BIOTECH-R1902: Fast-tracking the Development of BBTV-resistant Banana Cultivars through Modern Biotech Tools: Molecular Profiling towards Marker Development and Diagnostics (Phase I)” and Bill and Melinda Gates Foundation/University of Queensland’s project on “BBTV mitigation: Community management in Nigeria, and screening wild banana progenitors for resistance”. We gratefully acknowledge the Institute of Plant Breeding for the use of facilities and equipment. Also, the authors extend their warmest gratitude to Rodelio R. Pia, Ronilo M. Bajaro, Wilermie Driz-Hernandez, Reina S. Caro, Don Emanuel M. Cardona, Rowena B. Frankie, and Jen Daine L. Nocum for the technical and administrative support.

## Conflict of interest

The authors declare no conflict of interests.

## Notes

### Competing Interest Statement

The authors have declared no competing interest.

